# Estimating limit of detection of M. *tuberculosis* DNA as an analyte released from a chemically coated solid matrix and using a WHO-approved CB-NAAT platform

**DOI:** 10.1101/2023.09.26.558062

**Authors:** Krishna H. Goyani, Pratap N. Mukhopadhyaya

## Abstract

The limit of detection of Mtb (*Mycobacterium tuberculosis*) bacilli was evaluated by detection of released DNA from a chemically treated cellulose matrix. Loopful of the Mtb cells were suspended in phosphate buffer (pH 6.8), DNA extracted from it and quantified using a commercial Mtb real time PCR quantitation kit. Synthetic sputum, prepared using acrylamides as the matrix of choice was mixed with defined number of Mtb bacilli and spotted onto the cellulose matrix. Mtb DNA from the matrix was released into a DNA-release buffer and detected using a WHO-approved CB-NAAT detection platform. One hundred and fifty, 500 and 1000 but not 10 Mtb bacilli/mL of synthetic sputum could be detected by this protocol. The result underlined the potential of cellulose matrix as a potent, room temperature, Mtb-infected sputum transportation, storage and DNA-release device for sensitive detection of Mtb using *Mtb Xpert Ultra CB-NAAT* as the WHO-approved Mtb test of choice.

## Introduction

*Mycobacterium tuberculosis* (Mtb) is an ancient pathogen with more than 70,000 years of existence on earth. Its dramatic success in evolution is underlined by the fact that currently it is infecting nearly 1 billion people across the globe^1^. With the world recording around 10.4 million new TB infection cases every year, broad estimates indicate that almost one third of world’s population are likely carriers of this pathogen and hence carrying the risk of developing the disease^2^. The worry stems from the fact that the impact on a country’s exchequer is profound as the disease strikes people who are at the prime age of their lives^3^. Mtb diagnostics is evolving at a rapid pace as much as the therapeutics since the length of illness of this disease makes improved case-finding one of the crucial points for handling the disease^4^.

One of the most crucial aspects of initiating disease control for tuberculosis is rapid and accurate detection of the pathogen and also the determination of its profile towards resistance to drugs that are used to treat the disease^5^. In the area of culture-based diagnostics, several new developments have occurred. This includes for example, detection of growth at an early stage and stringent oxygen level control and employment of matrix-assisted laser desorption/ionization time-of-flight mass spectrometry (MALDI-TOF-MS) for identification of the pathogen^6^. However, it is worth mentioning that advent of nucleic acid amplification tests or NAATs had reduced the testing time compared to culture-based methods that take more than a week to generate results^7,8^. The NAATs primarily rely upon *in vitro* amplification of nucleic acids, either DNA or RNA and often involve techniques such as DNA sequencing and DNA-DNA hybridization to detect drug resistance^9^. Recently, the World Health Organization (WHO) has advocated development and use of sputum-based tests that can substitute smear microscopy and be used as an initial, rule-out test followed by a rapid drug sensitivity assay that can be done at a simple microscopy level laboratory^10^.

The world has seen several innovations in the domain of Mtb diagnostics. These include light-emitting diode microscope-based test, rapid automated liquid culture systems such as the Becton Dickinson MGIT 960, NAAT such as line probe assays and automated Cartridge based NAAT assays that include the Cepheid Xpert® MTB/RIF system (Cepheid, Inc., Sunnyvale, CA, USA)^11^.

The technique of DNA sequencing has shown that over 95% of all rifampicin resistant strains of Mtb harbour mutation with an 81-base pair (bp) stretch of the gene encoding for the beta subunit of RNA polymerase (rpoB)^12,13,14^. This laid the foundation for several rapid Mtb detection as well as drug resistant identification tests such as the GenoType MTBDRplus [Hain Lifescience GmbH, Nehren, Germany], INNO LIPA Rif.TB [Innogenetics, Ghent, Belgium]) and GeneXpert MTB/RIF; Cepheid, Sunnyvale, CA)^15,12,16^.

The GeneXpert MTB/RIF test is an integrated, cartridge-based Mtb diagnostic device that performs hand-free extraction of Mtb DNA from sputum followed by its detection by a hemi-nested real time PCR assay workflow. The test interrogates the 81 bp rpoB region and scores for detection of Mtb as well as its drug-resistance profile towards rifampicin^15,17^.

In this study, we attempted to estimate the limit of detection of Mtb DNA on this CB-NAAT platform using DNA released from a solid matrix-based sputum transportation and storage medium that was spotted with varying number of Mtb bacilli by mixing it with synthetic sputum formulated for this study.

## Materials and Methods

### Culture of Mtb bacilli

Three smear-positive clinical sputum specimens were decontaminated by treating with N-acetyl-L-cysteine and sodium hydroxide method (NALC-NaOH). Following centrifugation, the sediments were resuspended in 1.5 ml of sterile phosphate buffer (pH 6.8) and used to inoculate culture media. A smear prepared from the sediments were re-examined for the presence of acid-fast bacilli (AFB). For confirming the presence of Mtb, a liquid culture media based on fluorometric detection of growth was used. For this, 0.5 ml of each of the processed sputum samples were used to inoculate Mycobacteria Growth Indicator Tube (MGIT) tubes and incubated in the MGIT 960 instrument at 37°C. A volume of 0.25 ml of the processed sputum samples were also used to inoculate solid culture media (Lowenstein-Jensen or LJ; Hi Media, India) and incubated at 37°C. Once the tubes were provisionally identified as positive by growth, a smear of a sample from the tubes were prepared for examination for AFB (Acid-Fast Bacilli). All smears were stained by the Kinyoun method^18^ and examined with a light microscope. The Mtb strains isolated from the clinical specimens were identified by the MGIT TBc ID method (MPT 64: Becton Dickinson, Sparks, Maryland, USA). This is an immunochromatographic assay (ICA) that detects MPT64 antigen specifically secreted from Mtbc bacteria. Once the MTB complex strains were identified, drug susceptibility tests (DST) were performed using the MGIT SIRE (Becton Dickinson-Sparks, Maryland, USA) following the manufacturer’s instructions. For the test, the final concentration of 83 μg/ml of streptomycin (STR), 83μg/ml isoniazid (INH), 83 μg/ml rifampin (RIF) and 415 μg/ml ethambutol (EMB) were used.

All three specimens, labelled specimen 1, 2 and 3 were found to be true positive. The positive cultures were maintained in Lowenstein-Jensen (LJ) (Hi Media, India) and specimen no. 1 was selected for use in further downstream applications.

### Preparation of Mtb bacilli suspension

A loopful of Mtb culture (specimen no. 1) was gently taken from the LJ slant and suspended in 200 μL of phosphate buffer (pH 6.8; 37C). A volume of 100 μL was used for extraction of Mtb DNA while the remaining was stored at 4C for dilution and further processing.

### Estimating copy number of Mtb bacilli in the phosphate buffer suspension

To estimate copy number of the Mtb bacilli in phosphate buffer, 100 μL of the bacterial suspension was used to extract DNA and run a quantitative real time PCR reaction. DNA was extracted using a spin column-based Mtb DNA extraction kit (Wobble Base Bioresearch, India) and quantified using a HELINI *Mycobacterium tuberculosis* [MTB] Real-time PCR Kit (Helini Biomolecules, Chennai, India). Copy number of Mtb DNA in the suspension was computed using the formula as suggested by the manufacturer:

Result (copies/ml) = Result (copies/μL) x Elution Volume (μL) ÷ Sample Volume (ml).

All tests were run in triplicate and on a Rotor-Gene Q 2plex real time PCR Platform from QIAGEN (Germany).

### Preparing synthetic sputum with known quantity of Mtb bacilli

Synthetic sputum was prepared as described by Yamada et al., 2006^19^. Briefly, 22.2 grams of acrylamide was mixed with 0.6 grams of N, N’-methylenebisacrylamide and the volume made up to 100 ml with sterile distilled water. The reagent was labelled as the stock solution and stored at 4C in a sterile, amber-coloured bottle and protected from light. For preparing the synthetic sputum, 1.75 mL of the polyacrylamide stock was mixed with 6.25 mL sterile distilled water, 2 mL of Tris Borate EDTA (TGE) buffer (pH: 8.0) and 100 μL of 10% ammonium peroxidisulphate. To this, 8μL of a polymerization accelerator (TEMED) was added along with desired volume of Mtb suspension such that the final volume was 1 mL, and mixed thoroughly. It was then left at room temperature for 6 hours prior to use as Mtb-infected artificial sputum.

A test panel of five synthetic sputum suspensions were prepared in triplicate. Each of the 1 ml of synthetic sputum suspensions contained quantified number of the live Mtb pathogen cells and comprised of 10, 150, 500, 1000 & 10,000 CFU of Mtb bacilli respectively.

### Spotting a cellulose matrix with artificially infected synthetic human sputum

Mtb-infected synthetic sputum from the test panel were processed by mixing it with an equal volume of commercial Mtb cell lysis buffer solution (Wobble Base Bioresearch, India) and spotted onto a cellulose matrix soaked with chaotropic salts, surfactants, anti-fungal agents, a dye and heat-dried. Briefly, 1 ml of Mtb bacilli-containing synthetic sputum was added to an equal volume of the Mtb cell lysis buffer, mixed well by inverting the tube for 5-10 times and poured onto the cellulose matrix using a sterile disposable dropper and allowed to stand for a minimum period of 12 hours prior to capturing of the released Mtb DNA from the cellulose matrix.

*Release of Mtb DNA from TBSend card* The solid matrix was placed inside a wide mouth-bottle pre-filled with 3 ml of a sterile, isotonic buffer (DNA Release Buffer), vigorously shaken and incubated for 15 minutes. One ml of this processed buffer was used to run the Xpert Ultra CB-NAAT test for Mtb.

### Running the Mtb Xpert Ultra CB-NAAT test

The Xpert Ultra test was performed as described by the manufacturer. Briefly, 1 ml of the processed DNA-release buffer from each of the 4 specimens of the test panel (all replicates) were added to 1 ml of sample reagent (part of the Xpert Ultra test kit) and incubated for a period of 15 minutes. Two ml of this treated DNA-release buffer & sample reagent mixture was transferred to an Xpert Ultra cartridge and placed on the Xpert instrument to initiate the run. The test report was obtained after a period of around 80 minutes. As mentioned above, all samples of the test panel were processed in triplicate.

### Statistical analysis

All experiments were performed in triplicate and data are expressed as the mean ± standard deviation. The statistical analysis of differences was performed using the t-test of variance and Pearson r-test of Correlation with Graphpad prism 6.0 software and the P<0.05 was considered to indicate a statistically significant difference.

The study was approved by the Institutional Review Board of Nirmal Hospitals (Nirmal/HPL/Ethics/001) and performed in accordance with the principles of the Declaration of Helsinki.

## Results and Discussion

All 3 smear-positive sputum samples, *viz*., sputum specimen 1, 2 and 3 were found to be positive for Mtb infection upon further investigation. For this study, Mtb bacilli obtained from sputum specimen no. 1 was used for further investigations.

Mtb from the positive sputum sample (Sputum specimen 1) grew well in LJ slants and sufficient freshly-grown bacilli were available to prepare a visibly dense suspension in 200 μL of phosphate buffer.

DNA extracted from 100 μL of the phosphate buffer suspension-containing live Mtb bacilli were used to estimate the number of colony forming units (CFU) of Mtb at source. Real time quantitative PCR using a commercial kit indicated Mtb DNAcopy number of 8097.57 copies/μL in the phosphate buffer suspension (Figure 1) which was equivalent to 24,29,271 copies of CFU/mL. A volume of 0.41 μL, 6.17 μL, 20.58 μL, 41.16 μL and 410 μL, when mixed to prepare 1 mL of synthetic sputum, generated quantified 1 mL-synthetic sputum aliquots comprising of 10, 150, 500, 1000 and 10,000 Mtb bacilli/mL respectively. All suspensions were taken up for further study except 10,000 Mtb bacilli/ml vial which was subjected to microscopy by Ziehl-Neelsen staining ^20^.

**Figure 1:**
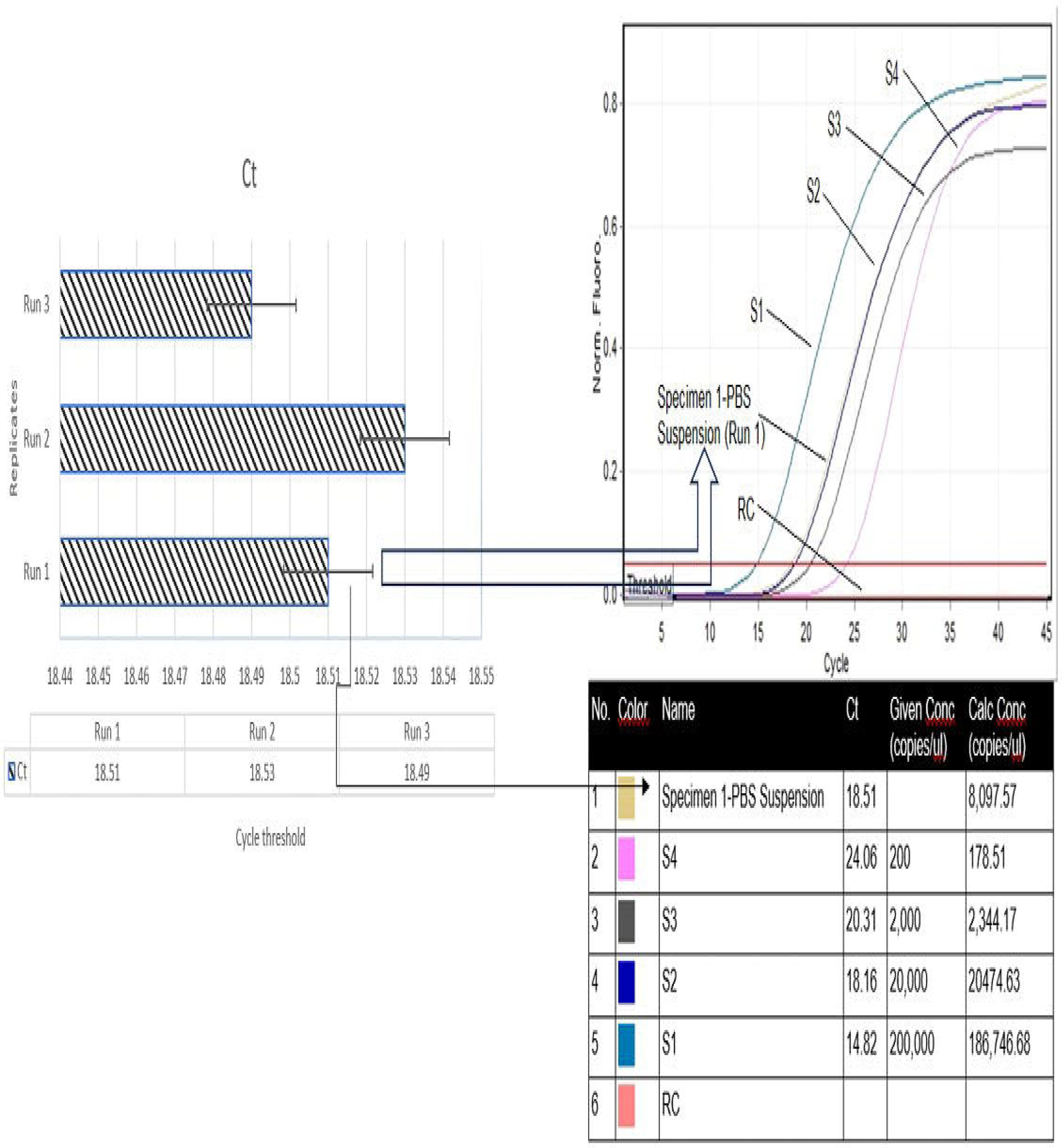
Real Tune PCR data of a replicate sample (Run 1) along with quantitation standards indicating computed Mtb bacilli DNA copy number (copies/μL) of the suspension prepared for generating the quantified, bacilli-embedded synthetic sputum panel and comprising of 10,150.500.1000 and 10.000 Mtb bacilli/mL.

Incubation of 15 minutes at room temperature along with intermittent shaking of the wide mouth bottle containing the Mtb bacilli-mixed synthetic sputum-spotted solid matrix suspended in 3 mL of DNA-release buffer effectively rescued trapped Mtb DNA into the solution.

The solid matrix is primarily of cellulose origin which is a hydroxylated polymer with enhanced affinity for DNA. In this study, it was impregnated with a range of chemicals including detergents, buffers, chelating and anti-fungal agents that further assisted in archiving and release of DNA when treated with appropriate and compatible buffer solution^21^.

One mL of this Mtb DNA-containing buffer was used to run Xpert Ultra cartridge-based Mtb NAAT. As per the prescribed protocol for running Xpert Ultra test, around 1 mL of sputum sample was added to an equal volume of sample reagent (a mixture of Sodium Hydroxide and Isopropanol) and incubated for 15 minutes prior to adding it into the cartridge for run.

The CB-NAAT could detect Mtb DNA from 1000, 500 and 150 but not the 10 Mtb bacilli/mL suspension-containing solid cellulose matrix (Table 1). All replicates generated equivalent data indicating the robustness of the assay (Supplementary data 1). Given the fact that detection of Mtb in the 10,000 bacilli/mL was obvious, the sample was not run on CB-NAAT platform. However, it was used to detect Mtb bacilli using the conventional sputum microscopy method^18^. The staining-based microscopy showed clear presence of elongated Mtb Bacilli, confirming the presence of pathogen cells in the synthetic sputum suspension (Figure 2).

**Table 1:** Cartridge-based nucleic acid amplification testing (Xpert® MTB/RIF Ultra; Cepheid’s GeneXpert® systems, USA) and stain-based microscopy results obtained from the Quantified Mtb bacilli-containing synthetic sputum suspensions.

| No. of *Mtb* bacilli/mL in synthetic sputum suspension | Xpert® MTB/RIF Ultra result | Staining-based microscopy result |
| --- | --- | --- |
| 10 | Not detected | Not processed |
| 150 | Detected, Low | Not processed |
| 500 | Detected, Low | Not processed |
| 1000 | Detected, Low | Not processed |
| 10,000 | Not run | Detected |

**Figure 2:**
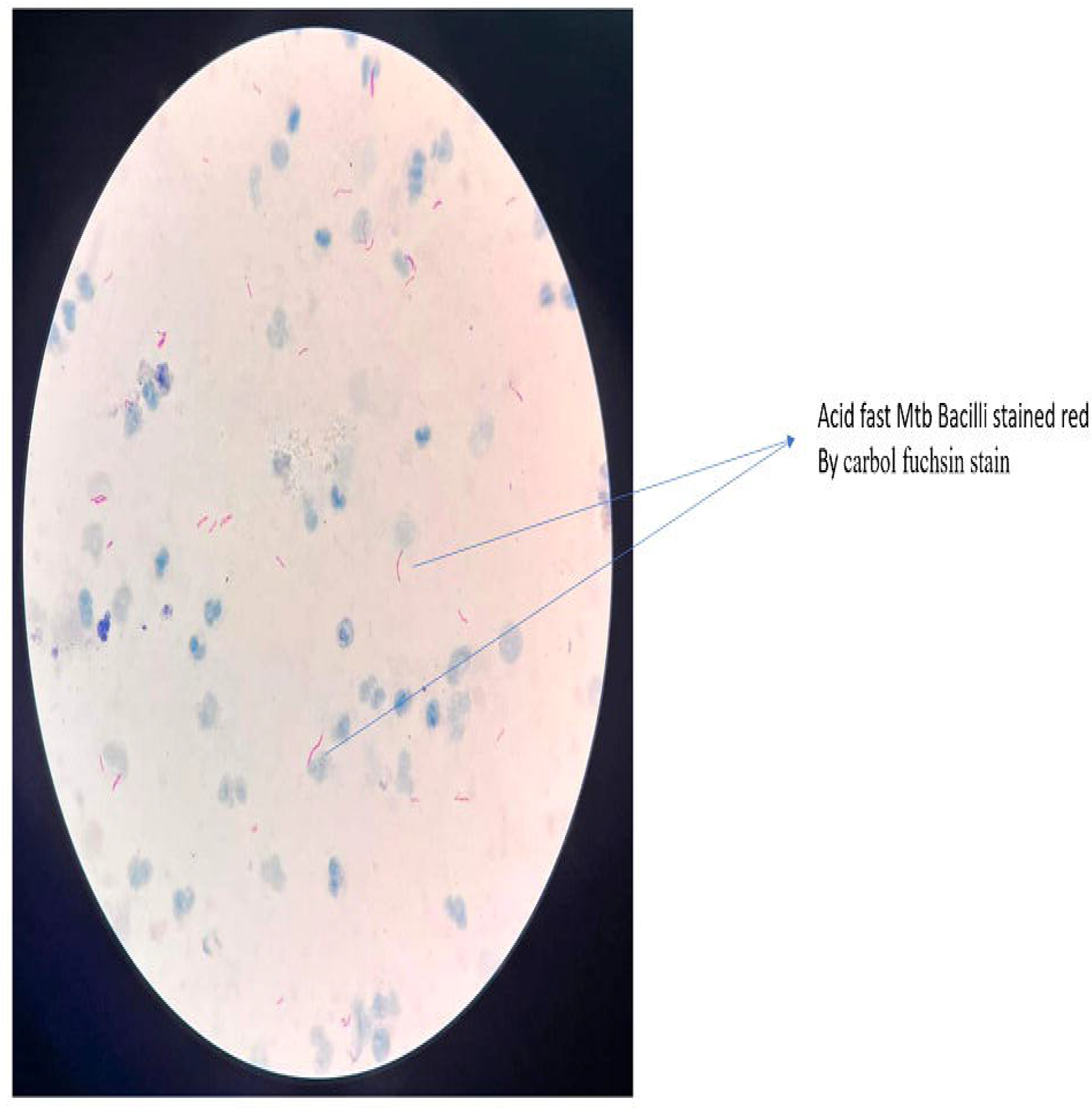
Photomicrograph of Mtb 10,000 bacilli/mL-suspension sample taken using a ZEISS Primo Star iLED microscope. Mtb bacilli were detected by die staining-based microscopy method using a carbol fuchsin stain, acid alcohol decolourizer, and methylene blue counterstain. Acid-fast Mtb bacilli were stained red and seen as elongated cells in this image (pointed arrows), while the background of debris stained blue (seen as dark patches of irregular size across the field). Hie data confirmed die presence of acid-fast mycobacteria in die suspension.

Around 10.4 million new tuberculosis infection cases occurred in the year 2015. However, as per record, just 6.1 million cases, amounting to 59% were successfully diagnosed^22^. In the same year, out of 5,80,000 cases of multidrug resistant tuberculosis cases, only 1,25,000 or 20% could be detected^21^.The primary reasons for this gap in occurrence and identification of the disease was absence of sufficiently sensitive and rapid diagnostics which was accessible to the population of a country in general^23^. This unmet need was successfully met by the Xpert MTB/RIF assay (Cepheid, Sunnyvale, CA, USA). The test is cartridge-based and because of the high-end technology used, it dramatically enhanced the detection rate of Mtb infection and improved identification of multi-drug resistant cases of tuberculosis^24,25,26^. Currently, more than 120 countries in the world use the Xpert MTB/RIF assay for detection of Mtb infection^27^. However, with time it was eventually observed that clinical samples with lesser number of Mtb bacilli could not be successfully detected by this test. The issue was more profound in smear-negative and extra pulmonary cases of Mtb infection where the number of Mtb bacilli are inherently low^28,29,30^. Such phenomena reduced the confidence on negative results generated by the kit and also inadvertently promoted empirical approach to tuberculosis management and overtreatment of patients^31,32^. In Mtb strains harbouring genetic mutations that are phenotypically silent, low bacterial load generated false positive results with Xpert MTB/RIF assay. Although a rare phenomenon, it reduced the diagnostic confidence on positive reports when the infection load was low^33,34^.

In this backdrop, the Xpert MTB/RIF Ultra assay (Xpert Ultra) was developed that effectively addressed the limitations of the earlier version of the test. With enhanced assay chemistry, improved cartridge design and simultaneous detection of two different multi copy targets (IS6110 & IS1081), the limit of detection and other diagnostic features of this version of Xpert MTB assay was superior and more effective^35^. Amongst other advantages extended by this new version of the Xpert Mtb detection assay, a significant one was improved limit of detection by 1-log value compared to its predecessor^35^.

The cellulose matrix used in this study has potential to circumvent several disadvantages associated with traditional method of transportation and storage of sputum for NAAT. This is possible due to the use of a this chemically coated cellulose matrix that has the property to lyse Mtb bacilli cells, trap its DNA, store at room temperature and finally release it on demand after an extended period of time (up to 6 years). This solid matrix, with its cent per cent bactericidal properties, has excellent biosafety features also (data not shown) and potential to be a tool of choice for sputum transportation, storge and DNA extraction of Mtb for NAAT.

This study was intended to establish the clinical sensitivity of the chemically coated cellulose matrix when Xpert MTB/RIF Ultra assay was the test of choice for detection of Mtb infection.

One hundred and fifty Mtb bacilli per mL was successfully detected by the Xpert MTB/RIF Ultra assay system when the cellulose matrix was the medium of choice for room temperature storage and release of Mtb DNA. All triplicate samples generated identical data thus enhancing the confidence on the observation. As expected, apart from 150 bacilli/mL of artificial sputum samples, 500 and 1000 Mtb bacilli per 1 mL samples were also detected in all replicates. However, the group of the synthetic sputum samples (n=3) loaded with 10 Mtb bacilli per mL could not be detected by the Xpert MTB/RIF Ultra assay in all triplicates. Given the fact that Xpert MTB/RIF Ultra assay has a clinical sensitivity of 15.6 CFU/mL^34^, this result was expected.

## Conclusion

The results from this study indicated that cellulose matrixes can be successfully and effectively used as a transportation, storage and DNA-release device for Mtb sputum to detect Mtb bacilli with a limit of detection of 150 bacilli per mL of sputum or lower, but not 10 bacilli/mL, when Xpert MTB/RIF Ultra assay was the test of choice of detection of Mtb.

## Acknowledgements

This study was supported by Grand Challenge-TB Control (GCTBC) grants, namely GCTBC/C2P1/2015/09/30/01 & GCTBC/C2P2/2017/04/01/01 and funded by Industry Research Assistance Council (BIRAC) of the Government of India, United States Agency for International Development (USAID), and Bill & Melinda Gates Foundation. IKP Knowledge Park (Hyderabad, India) was the implementation partner for these programmes.

## Notes

### Competing Interest Statement

The authors have declared no competing interest.

## References

1. MacDonald, E. M., & Izzo, A. A. (2015). Tuberculosis Vaccine Development. In: Ribbon W (Ed.). Tuberculosis-expanding knowledge. In Tech.

2. Global Tuberculosis Report 2016. World Health Organization. Available at: http://www.who.int/tb/publications/global_repor t/en/[Accessed on 06/12/2016].

3. Dye C. (2006). Global epidemiology of tuberculosis. Lancet (London, England), 367(9514), 938–940.

4. Iseman M. D. (2000). TB elimination in the 21st century, a quixotic dream?. The international journal of tuberculosis and lung disease : the official journal of the International Union against Tuberculosis and Lung Disease, 4(12 Suppl 2), S109–S110.

5. Engström A. (2016). Fighting an old disease with modern tools: characteristics and molecular detection methods of drug-resistant Mycobacterium tuberculosis. Infectious diseases (London, England), 48(1), 1–17.

6. Ghodbane, R., Raoult, D., & Drancourt, M. (2014). Dramatic reduction of culture time of Mycobacterium tuberculosis. Scientific reports, 4, 4236.

7. Cho, W. H., Won, E. J., Choi, H. J., Kee, S. J., Shin, J. H., Ryang, D. W., & Suh, S. P. (2015). Comparison of AdvanSure TB/NTM PCR and COBAS TaqMan MTB PCR for Detection of Mycobacterium tuberculosis Complex in Routine Clinical Practice. Annals of laboratory medicine, 35(3), 356–361.

8. Huggett, J. F., McHugh, T. D., & Zumla, A. (2003). Tuberculosis: amplification-based clinical diagnostic techniques. The international journal of biochemistry & cell biology, 35(10), 1407–1412.

9. Balasingham, S. V., Davidsen, T., Szpinda, I., Frye, S. A., & Tønjum, T. (2009). Molecular diagnostics in tuberculosis: basis and implications for therapy. Molecular diagnosis & therapy, 13(3), 137–151.

10. Denkinger, C. M., Kik, S. V., Cirillo, D. M., Casenghi, M., Shinnick, T., Weyer, K., Gilpin, C., Boehme, C. C., Schito, M., Kimerling, M., & Pai, M. (2015). Defining the needs for next generation assays for tuberculosis. The Journal of infectious diseases, 211 Suppl 2(Suppl 2), S29–S38.

11. Drobniewski, F., Nikolayevskyy, V., Maxeiner, H., Balabanova, Y., Coppola, N., Kontsevaya, I., & Ignatyeva, O. (2013). Rapid diagnostics of tuberculosis and drug resistance in the industrialized world: clinical and public health benefits and barriers to implementation. BMC Medicine, 11:190.

12. Boehme, C. C., Nicol, M. P., Nabeta, P., Michael, J. S., Gotuzzo, E., Tahirli, R., Gler, M. T., Blakemore, R., Worodria, W., Gray, C., Huang, L., Caceres, T., Mehdiyev, R., Raymond, L., Whitelaw, A., Sagadevan, K., Alexander, H., Albert, H., Cobelens, F., Cox, H., … Perkins, M. D. (2011). Feasibility, diagnostic accuracy, and effectiveness of decentralised use of the Xpert MTB/RIF test for diagnosis of tuberculosis and multidrug resistance: a multicentre implementation study. Lancet (London, England), 377(9776), 1495–1505.

13. Mani, C., Selvakumar, N., Narayanan, S., & Narayanan, P. R. (2001). Mutations in the rpoB gene of multidrug-resistant Mycobacterium tuberculosis clinical isolates from India. Journal of clinical microbiology, 39(8), 2987–2990.

14. Telenti, A., Imboden, P., Marchesi, F., Lowrie, D., Cole, S., Colston, M. J., Matter, L., Schopfer, K., & Bodmer, T. (1993). Detection of rifampicin-resistance mutations in Mycobacterium tuberculosis. Lancet (London, England), 341(8846), 647–650.

15. Blakemore, R., Story, E., Helb, D., Kop, J., Banada, P., Owens, M. R., Chakravorty, S., Jones, M., & Alland, D. (2010). Evaluation of the analytical performance of the Xpert MTB/RIF assay. Journal of clinical microbiology, 48(7), 2495–2501.

16. Cirillo, D. M., Piana, F., Frisicale, L., Quaranta, M., Riccabone, A., Penati, V., Vaccarino, P., & Marchiaro, G. (2004). Direct rapid diagnosis of rifampicin-resistant M. tuberculosis infection in clinical samples by line probe assay (INNO LiPA Rif-TB). The new microbiologica, 27(3), 221–227.

17. Helb, D., Jones, M., Story, E., Boehme, C., Wallace, E., Ho, K., Kop, J., Owens, M. R., Rodgers, R., Banada, P., Safi, H., Blakemore, R., Lan, N. T., Jones-López, E. C., Levi, M., Burday, M., Ayakaka, I., Mugerwa, R. D., McMillan, B., Winn-Deen, E., … Alland, D. (2010). Rapid detection of Mycobacterium tuberculosis and rifampin resistance by use of on-demand, near-patient technology. Journal of clinical microbiology, 48(1), 229–237.

18. Salleh, F. M., Al-Mekhlafi, A. M., Nordin, A., Yasin, M., Al-Mekhlafi, H. M., & Moktar, N. (2011). Evaluation of gram-chromotrope kinyoun staining technique: its effectiveness in detecting microsporidial spores in fecal specimens. Diagnostic microbiology and infectious disease, 69(1), 82–85.

19. Yamada, H., Mitarai, S., Aguiman, L., Matsumoto, H., & Fujiki, A. (2006). Preparation of mycobacteria-containing artificial sputum for TB panel testing and microscopy of sputum smears. The international journal of tuberculosis and lung disease : the official journal of the International Union against Tuberculosis and Lung Disease, 10(8), 899–905.

20. Laifangbam, S., Singh, H. L., Singh, N. B., Devi, K. M., & Singh, N. T. (2009). A comparative study of fluorescent microscopy with Ziehl-Neelsen staining and culture for the diagnosis of pulmonary tuberculosis. Kathmandu University medical journal (KUMJ), 7(27), 226–230.

21. Preetha J Shetty (2020). The Evolution of DNA Extraction Methods.AJBSR.MS.ID.001234, 8(1).

22. WHO (2016). Global tuberculosis report 2016. Geneva: World Health Organization, 2016.

23. Uplekar, M., Weil, D., Lonnroth, K., Jaramillo, E., Lienhardt, C., Dias, H. M., Falzon, D., Floyd, K., Gargioni, G., Getahun, H., Gilpin, C., Glaziou, P., Grzemska, M., Mirzayev, F., Nakatani, H., Raviglione, M., & for WHO’s Global TB Programme (2015). WHO’s new end TB strategy. Lancet (London, England), 385(9979), 1799–1801.

24. Boehme, C. C., Nabeta, P., Hillemann, D., Nicol, M. P., Shenai, S., Krapp, F., Allen, J., Tahirli, R., Blakemore, R., Rustomjee, R., Milovic, A., Jones, M., O’Brien, S. M., Persing, D. H., Ruesch-Gerdes, S., Gotuzzo, E., Rodrigues, C., Alland, D., & Perkins, M. D. (2010). Rapid molecular detection of tuberculosis and rifampin resistance. The New England journal of medicine, 363(11), 1005–1015.

25. Steingart, K. R., Schiller, I., Horne, D. J., Pai, M., Boehme, C. C., & Dendukuri, N. (2014). Xpert® MTB/RIF assay for pulmonary tuberculosis and rifampicin resistance in adults. The Cochrane database of systematic reviews, 2014(1), CD009593.

26. World Health Organization. (2013). Xpert MTB/RIF assay for the diagnosis of pulmonary and extrapulmonary TB in adults and children: policy update. Geneva: World Health Organization, 2013.

27. World Health Organization. (2014). WHO monitoring of Xpert MTB/RIF roll-out. Geneva: World Health Organization, 2014.

28. Nicol, M. P., Workman, L., Isaacs, W., Munro, J., Black, F., Eley, B., Boehme, C. C., Zemanay, W., & Zar, H. J. (2011). Accuracy of the Xpert MTB/RIF test for the diagnosis of pulmonary tuberculosis in children admitted to hospital in Cape Town, South Africa: a descriptive study. The Lancet. Infectious diseases, 11(11), 819–824.

29. Theron, G., Peter, J., van Zyl-Smit, R., Mishra, H., Streicher, E., Murray, S., Dawson, R., Whitelaw, A., Hoelscher, M., Sharma, S., Pai, M., Warren, R., & Dheda, K. (2011). Evaluation of the Xpert MTB/RIF assay for the diagnosis of pulmonary tuberculosis in a high HIV prevalence setting. American journal of respiratory and critical care medicine, 184(1), 132–140.

30. Sohn, H., Aero, A. D., Menzies, D., Behr, M., Schwartzman, K., Alvarez, G. G., Dan, A., McIntosh, F., Pai, M., & Denkinger, C. M. (2014). Xpert MTB/RIF testing in a low tuberculosis incidence, high-resource setting: limitations in accuracy and clinical impact. Clinical infectious diseases: an official publication of the Infectious Diseases Society of America, 58(7), 970–976.

31. Theron, G., Peter, J., Dowdy, D., Langley, I., Squire, S. B., & Dheda, K. (2014). Do high rates of empirical treatment undermine the potential effect of new diagnostic tests for tuberculosis in high-burden settings? The Lancet. Infectious diseases, 14(6), 527–532.

32. Theron, G., Zijenah, L., Chanda, D., Clowes, P., Rachow, A., Lesosky, M., Bara, W., Mungofa, S., Pai, M., Hoelscher, M., Dowdy, D., Pym, A., Mwaba, P., Mason, P., Peter, J., Dheda, K., & TB-NEAT team (2014). Feasibility, accuracy, and clinical effect of point-of-care Xpert MTB/RIF testing for tuberculosis in primary-care settings in Africa: a multicentre, randomised, controlled trial. Lancet (London, England), 383(9915), 424–435.

33. Mathys, V., van de Vyvere, M., de Droogh, E., Soetaert, K., & Groenen, G. (2014). False-positive rifampicin resistance on Xpert® MTB/RIF caused by a silent mutation in the rpoB gene. The international journal of tuberculosis and lung disease : the official journal of the International Union against Tuberculosis and Lung Disease, 18(10), 1255–1257.

34. Ocheretina, O., Byrt, E., Mabou, M. M., Royal-Mardi, G., Merveille, Y. M., Rouzier, V., Fitzgerald, D. W., & Pape, J. W. (2016). Falsepositive rifampin resistant results with Xpert MTB/RIF version 4 assay in clinical samples with a low bacterial load. Diagnostic microbiology and infectious disease, 85(1), 53–55.

35. Chakravorty, S., Simmons, A. M., Rowneki, M., Parmar, H., Cao, Y., Ryan, J., Banada, P. P., Deshpande, S., Shenai, S., Gall, A., Glass, J., Krieswirth, B., Schumacher, S. G., Nabeta, P., Tukvadze, N., Rodrigues, C., Skrahina, A., Tagliani, E., Cirillo, D. M., Davidow, A., … Alland, D. (2017). The New Xpert MTB/RIF Ultra: Improving Detection of Mycobacterium tuberculosis and Resistance to Rifampin in an Assay Suitable for Point-of-Care Testing. mBio, 8(4), e00812–17.

