## Supplementary data 1 for "Estimating limit of detection of M. *tuberculosis* DNA as an analyte released from a chemically coated solid matrix and using a WHO-approved CB-NAAT platform"

Patient ID: Sputum-150
Sample ID*: 309100454
Test Type: Specimen
Sample Type: Sputum

Assay Information

| Assay | Assay Version | Assay Type |
|---|---|---|
| Xpert MTB-RIF Ultra | 4 | In Vitro Diagnostic |

Test Result: MTB DETECTED LOW;
RIF Resistance DETECTED

Analyte Result

| Analyte Name | Ct | EndPt | Analyte Result | Probe Check Result |
|---|---|---|---|---|
| SPC | 24.1 | 144 | NA | PASS |
| IS1081-IS6110 | 20.3 | 507 | NA | PASS |
| rpoB1 | 23.1 | 404 | POS | PASS |
| rpoB2 | 25.6 | 143 | POS | PASS |
| rpoB3 | 24.4 | 200 | POS | PASS |
| rpoB4 | 26.0 | 125 | POS | PASS |

| | | | |
|---|---|---|---|
| User: | <None> | | |
| Status: | Done | Start Time: | 06/09/23 5:57:50 PM |
| Expiration Date*: | 28/07/24 | End Time: | 06/09/23 7:16:25 PM |
| S/W Version: | 6.4 | Instrument S/N: | 110000827 |
| Cartridge S/N*: | 1206657833 | Module S/N: | 671651 |
| Reagent Lot ID*: | 39207 | Module Name: | B1 |
| Notes: | | | |
| Error Status: | OK | | |

Errors
<None>

For In Vitro Diagnostic Use Only.

A

Patient ID: Sputum-500
Sample ID*: 309100453
Test Type: Specimen
Sample Type: Sputum

Assay Information

| Assay | Assay Version | Assay Type |
|---|---|---|
| Xpert MTB-RIF Ultra | 4 | In Vitro Diagnostic |

Test Result: MTB DETECTED LOW;
RIF Resistance DETECTED

Analyte Result

| Analyte Name | Ct | EndPt | Analyte Result | Probe Check Result |
|---|---|---|---|---|
| SPC | 24.9 | 135 | NA | PASS |
| IS1081-IS6110 | 21.1 | 468 | NA | PASS |
| rpoB1 | 24.0 | 361 | POS | PASS |
| rpoB2 | 27.4 | 121 | POS | PASS |
| rpoB3 | 25.5 | 176 | POS | PASS |
| rpoB4 | 28.4 | 101 | POS | PASS |

| | | | |
|---|---|---|---|
| User: | <None> | | |
| Status: | Done | Start Time: | 06/09/23 4:57:44 PM |
| Expiration Date*: | 28/07/24 | End Time: | 06/09/23 6:15:28 PM |
| S/W Version: | 6.4 | Instrument S/N: | 110000827 |
| Cartridge S/N*: | 1206657835 | Module S/N: | 659545 |
| Reagent Lot ID*: | 39207 | Module Name: | B4 |
| Notes: | | | |
| Error Status: | OK | | |

Errors
<None>

For In Vitro Diagnostic Use Only.

B

| | |
|---|---|
| Patient ID: | Suputum-1000 |
| Sample ID*: | 309100452 |
| Test Type: | Specimen |
| Sample Type: | Sputum |

Assay Information

| Assay | Assay Version | Assay Type |
|---|---|---|
| Xpert MTB-RIF Ultra | 4 | In Vitro Diagnostic |

Test Result: MTB DETECTED LOW;
RIF Resistance DETECTED

Analyte Result

| Analyte Name | Ct | EndPt | Analyte Result | Probe Check Result |
|---|---|---|---|---|
| SPC | 25.1 | 135 | NA | PASS |
| IS1081-IS6110 | 21.2 | 482 | NA | PASS |
| rpoB1 | 24.2 | 386 | POS | PASS |
| rpoB2 | 26.7 | 134 | POS | PASS |
| rpoB3 | 25.2 | 200 | POS | PASS |
| rpoB4 | 27.2 | 119 | POS | PASS |

| | | | |
|---|---|---|---|
| User: | <None> | | |
| Status: | Done | Start Time: | 06/09/23 4:36:45 PM |
| Expiration Date*: | 28/07/24 | End Time: | 06/09/23 5:55:26 PM |
| S/W Version: | 6.4 | Instrument S/N: | 110000827 |
| Cartridge S/N*: | 1206657877 | Module S/N: | 671651 |
| Reagent Lot ID*: | 39207 | Module Name: | B1 |
| Notes: | | | |
| Error Status: | OK | | |

Errors
<None>

For In Vitro Diagnostic Use Only.

**C**

**Supplementary data:** A, B & C: Xpert MTB Rif Ultra report of three samples of the Mtb-infected synthetic sputum panel, namely 150, 500 and 1000 indicating "MTB DETECTED" status. The semi quantitative value indicated in all three samples show "Low" and further, the drug resistance pattern deduced due to detection of specific mutation within the rpoB gene within the Mtb genome is identical in all the three reports since the Mtb bacilli originated from a single source in all the three samples.
